# A Systematic Review and Independent Benchmarking of Automated Nerve Morphometry Methods

**DOI:** 10.64898/2026.06.15.731646

**Authors:** Benton Chuter, Min Young Kim, Andrew B. Stiemke, Nikhil Dave, Harrison R. Cape, Zirui Zhou, Jay Herrin, Matthew Miller, William White, T.J. Hollingsworth, Monica M. Jablonski

**Affiliations:** Department of Ophthalmology, Hamilton Eye Institute, The University of Tennessee Health Science Center, Memphis, TN 38163, USA; Department of Genetics, Genomics and Informatics, The University of Tennessee Health Science Center, Memphis, TN 38163, USA; College of Medicine, University of Tennessee Health Science Center, Memphis, TN 38163, USA; Frederick P. Whiddon College of Medicine, University of South Alabama, Mobile, AL 36617, USA

**Author notes:** **Corresponding Author:** Monica M. Jablonski, PhD, FARVO Hamilton Eye Institute, 930 Madison Avenue, Memphis, TN 38163.

**Keywords:** Optic nerve, Morphometrics, Computer Vision, Deep Learning, Automated

## Abstract

**Objective:** To systematically review automated nerve morphometry tools and independently benchmark their performance on independent optic nerve datasets.

**Design:** Systematic review and comparative benchmarking study.

**Controls:** Benchmarking was performed using paraphenylenediamine-stained mouse (n = 85) and rat (n = 44) optic nerve images with manually annotated axon counts as ground truth.

**Methods:** Published studies describing automated or semi-automated neural tissue morphometry tools were identified through systematic searches of PubMed, Embase, and Scopus through January 2026 following PRISMA guidelines. Data extraction covered 70 fields across tool capabilities, imaging modality, species, automation level, and validation approach. Eighteen eligible tools (8 deep learning [DL], 10 classical computer vision [CV]) were benchmarked on both mouse and rat independent datasets.

**Main Outcome Measures:** Performance was assessed by mean absolute percentage error (MAPE), Pearson correlation, and median predicted-to-ground-truth ratio. Tools were ranked per image and compared using Friedman tests with Nemenyi post-hoc analysis.

**Results:** Seventy-one studies met inclusion criteria, spanning from 1999 to 2026. Deep learning methods represented 38% (27/71) of studies, increasing from 0% before 2017 to over 55% of publications after 2020. Axon counting was the most common output (73%, 52/71), while only 35% (25/71) reported g-ratio. Among benchmarked tools, Marina (CV, 2010) achieved the lowest average MAPE (32.9%). The top five tools (MAPE ranging from 32.9 to 44.8%) included both CV and DL methods and were statistically indistinguishable by Friedman-Nemenyi analysis (p > 0.05). Performance varied substantially across datasets: AxonJ (CV) achieved the second best MAPE on rat images (27.7%) but the worst on mouse images (438.6%).

**Conclusions:** No single tool demonstrated consistently superior performance across both datasets. Classical and deep learning approaches achieved comparable accuracy for axon counting. Tool selection should be guided by target species, tissue preparation protocol, and desired morphometric outputs. This systematic review and independent benchmarking study provide an evidence base for tool selection in optic nerve research.

## Introduction

Optic nerve degeneration is the defining pathological feature of glaucoma, the leading cause of irreversible blindness worldwide.^1,2^ Quantification of retinal ganglion cell (RGC) axons in optic nerve cross-sections remains the standard endpoint for evaluating neuroprotective therapies in preclinical models.^3,4^ Manual axon counting is labor-intensive, subjective, and impractical for large-scale studies, which has driven the development of automated morphometry tools over the past two decades.^5,6^ Early approaches relied on classical computer vision (CV) techniques such as thresholding, edge detection, and watershed segmentation,^7,8^ whereas more recent methods increasingly employ deep learning (DL), particularly convolutional neural networks trained on manually annotated datasets.^9,10^ Both approaches have been applied to optic nerve, sciatic nerve, spinal cord, and other neural tissues across species ranging from rodents to humans.^11,12^

Despite the growing number of available tools, selecting an appropriate morphometry method remains challenging. Published validations are typically performed on a single dataset collected by the tool developers, making cross-tool comparison difficult.^13,14^ Tools vary widely in their output capabilities; some provide only axon counts, while others measure fiber diameter, myelin thickness, g-ratio, and size distributions. Availability of source code, graphical user interfaces (GUIs), and documentation also varies.^15^ To date, no prior study has systematically cataloged these tools or independently compared their performance on shared datasets with consistent ground truth annotations.

To address these gaps, we conducted a systematic review of automated and semi-automated neural tissue morphometry tools, followed by an independent benchmarking study of optic nerve axon counting performance. We cataloged tool capabilities, imaging requirements, and validation approaches reported in the literature and benchmarked eligible tools on two optic nerve datasets with manual ground-truth counts. Tools were compared using standardized error metrics and nonparametric rank-based statistical tests. Our goal was to provide an evidence base to guide tool selection for optic nerve morphometry research.

## Methods

### Systematic Review Protocol

This systematic review was conducted following the Preferred Reporting Items for Systematic Reviews and Meta-Analyses (PRISMA) 2020 guidelines.^16^ The protocol was developed prospectively (B.C.) using search strategies detailed in **Supplementary Table 1.** We searched PubMed, Embase, and Scopus from database inception through January 2026. The search combined terms across four concept domains: (1) neural tissue types (optic nerve, axon, myelin, retinal ganglion cell, white matter, glia); (2) quantification methods (morphometry, axon counting, segmentation, grading, measurement); (3) computational approaches (automated, image analysis, digital pathology, machine learning, deep learning, convolutional neural network (CNN), ImageJ, watershed, thresholding); and (4) histological context (histology, histopathology, microscopy, immunohistochemistry). Reference lists of included studies were also screened for additional eligible publications.

### Eligibility Criteria

Studies were included if they described original research involving automated or semi-automated computational tools for quantification, segmentation, or morphometric analysis of neural tissue histology. The target population included human or animal optic nerve tissue, retinal ganglion cells, axons, myelin, and associated glial cells. Studies were required to report quantitative outcomes such as axon or myelin counts, morphometric measurements, segmentation accuracy, or detection performance metrics. Only peer-reviewed original research articles published in English were included. Reviews, editorials, conference abstracts without full data, and non-peer-reviewed reports were excluded. Complete inclusion and exclusion criteria are provided in **Supplementary Table S2.**

### Study Selection and Data Extraction

Two reviewers (B.C. and Z.Z.) independently screened titles and abstracts, followed by full-text review of potentially eligible studies. Disagreements were resolved by consensus with a third reviewer (N.D.). Data were extracted and verified (J.H., M.M. and M.Y.K.) into a standardized form covering 71 fields, including: study identifiers (authors, year, journal), tool characteristics (name, algorithm type, open-source availability, GUI), imaging parameters (modality, staining, magnification, resolution), biological context (species, nerve type, tissue state), validation approach, and 10 morphometric output categories (axon count, fiber count, axon diameter, fiber diameter, myelin thickness, g-ratio [ratio of axon diameter to the total fiber diameter],^17^ axon area, myelin area, fiber density, and size distribution).

### Algorithm Classification

Tools were classified into two categories based on their primary algorithmic approach. “Deep Learning” included tools using convolutional neural networks, U-Net architectures, or other deep learning frameworks as the primary analytical component. “Classical CV/Other” encompassed all other computational methods, including thresholding, edge detection, watershed segmentation, stereology, random forest classifiers, and other machine learning approaches that do not use deep neural networks. When a tool combined multiple approaches (e.g., deep learning with classical post-processing), it was classified by its primary segmentation or detection method.

### Independent Benchmarking

#### Tool Selection

From the pool of tools identified in the systematic review, 18 were selected for benchmarking based on the following criteria: (1) the tool must perform axon counting and/or myelin segmentation (the most common morphometric output); (2) source code or executable software must be accessible; and (3) the tool must be able to process standard 2D histological brightfield microscope images. Tools were identified by their reported software or algorithm name when available. For studies that did not assign a formal tool name, tools were referenced by the first author of the original publication for consistency throughout the review and benchmarking analysis. The 18 tools comprised 8 deep learning methods (AxonDeepSeg [ADS] Generalist, ADS Bright-Field Light,^9^ AxoNet,^13^ AxoNet 2.0,^14^ ls_axon_segmentation,^18^ Hifuku,^19^ Moiseev,^20^ and AMND^21^) and 10 classical CV methods (Marina,^22^ Reynaud APP,^5^ QuPath,^23^ g- ratio,^24^ Axoquant 2.0,^25^ Engelmann,^12^ Hessian/McCreedy,^26^ AxonSeg,^17^ AxonCounter,^27^ and AxonJ^27^).

#### Datasets

Two independent optic nerve datasets with manual ground truth axon counts were used. The mouse dataset consisted of 75 paraphenylenediamine (PPD)-stained archival mouse optic nerve cross-section images from mice across multiple age groups and experimental conditions. The heterogenous stock rat dataset consisted of 44 PPD-stained rat optic nerve cross-section images. For each image, ground truth axon counts were obtained through manual annotation by trained observers as described in our previous studies.^28,29^ These two datasets provided complementary test conditions: different species (mouse vs. rat), ranges of axon density, and tissue quality.

#### Performance Metrics

Each tool was applied to all images in both datasets. Tool outputs were matched to ground truth counts on a per-image basis. The following metrics were computed: mean absolute percentage error (MAPE); calculated as the mean of |predicted - ground truth| / ground truth across all images; Pearson correlation coefficient (r) between predicted and ground truth counts; mean absolute error (MAE); root mean square error (RMSE); and median ratio of predicted to ground truth counts. N_matched denotes the number of images for which the tool produced a valid, nonzero output that could be compared against ground truth.

MAPE was selected as the primary accuracy metric because it normalizes errors relative to the ground truth count, allowing for comparison across images with different axon densities. MAE quantifies the average magnitude of counting error in absolute error counts, whereas RMSE places greater weight on large errors to account for large misestimates. Pearson *r* assesses the strength of the linear relationship between predicted and ground truth counts but does not measure absolute agreement. The median predicted to ground truth ratio was used to identify systematic overcounting or undercounting bias, where a value of 1.0 indicates unbiased prediction, values >1.0 indicate overcounting, and value indicate <1.0 undercounting.

For tools producing pixel-level segmentation masks, segmentation accuracy was evaluated using the Dice similarity coefficient. Dice is a measure of spatial overlap between predicted and ground truth annotation ranging from 0 (no overlap) to 1 (complete overlap).^30^

### Statistical Analysis

Tools were ranked per image based on absolute percentage error. The Friedman test was used to assess whether tool rankings differed across images, with Nemenyi post-hoc tests to identify specific pairwise differences.^31^ Critical difference diagrams were generated for: (a) combined multi-metric rankings across both datasets, (b) the mouse dataset alone, and (c) the rat dataset alone. Statistical significance was set at p < 0.05.

## Results

### Systematic Review

#### Study Selection

The database search yielded 3951 records after deduplication. Following title and abstract screening, 91 studies underwent full-text review. Seventy-one studies met all inclusion criteria and were included in the final analysis. The complete PRISMA flow diagram with screening numbers is shown in **Figure 1**. The extracted dataset is provided in **Supplementary Table S3**.

**Figure 1.**
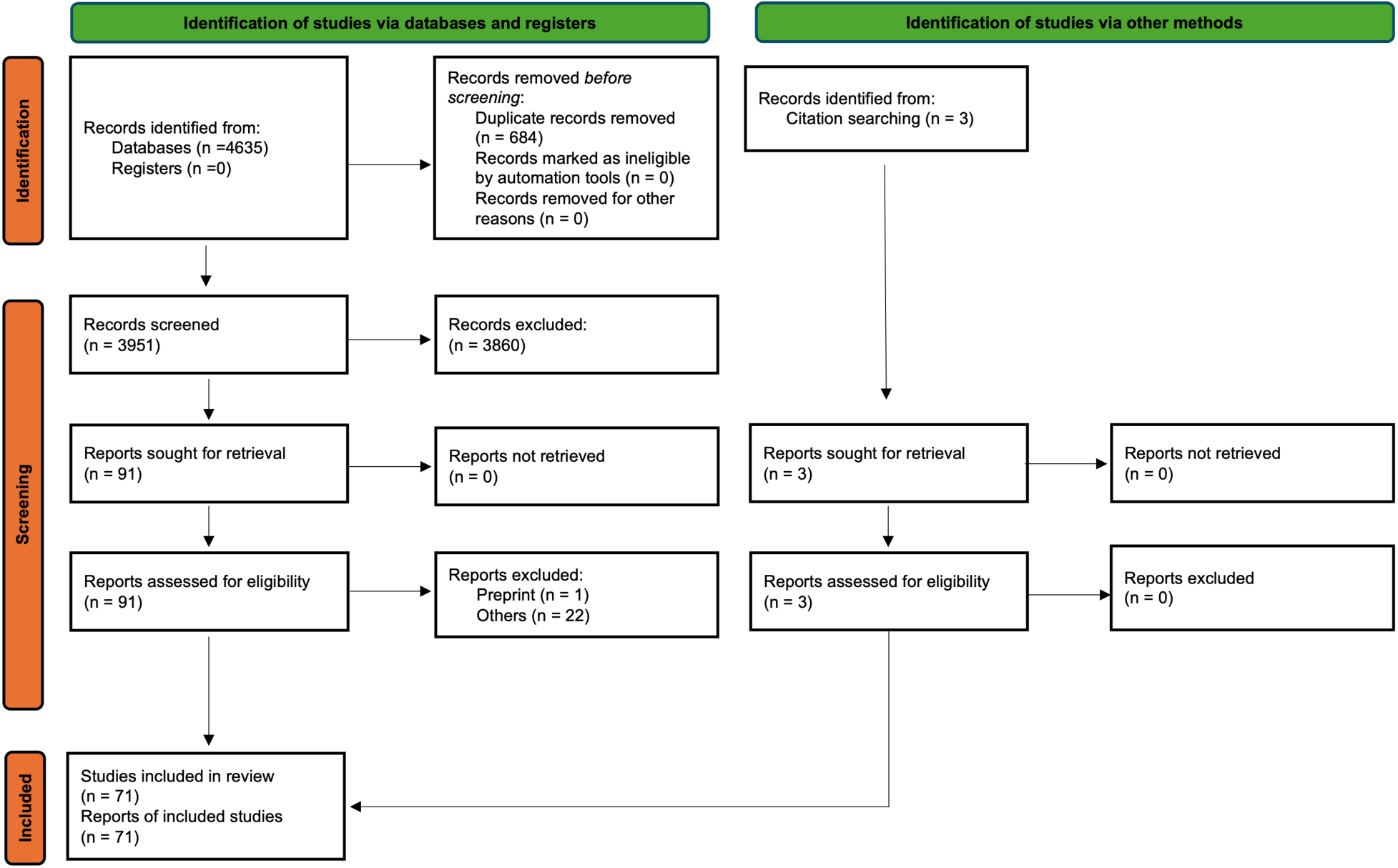
PRISMA 2020 flow diagram for the systematic review.^16^ Of 3951 unique records screened across PubMed, EMBASE, and Scopus, 91 underwent full-text review, and 71 manuscripts describing semi-automated and fully automated axon and/or myelin segmentation tools met inclusion criteria. Per-stage exclusion reasons are shown in the diagram.

#### Study Characteristics

The 71 included studies spanned 1999 to 2026 (**Table 1**). Light microscopy was the most common imaging modality (39%, 28/71), followed by transmission electron microscopy (TEM; 18%, 13/71), fluorescence microscopy (18%, 13/71), and confocal microscopy (13%, 9/71).

**Table 1.**
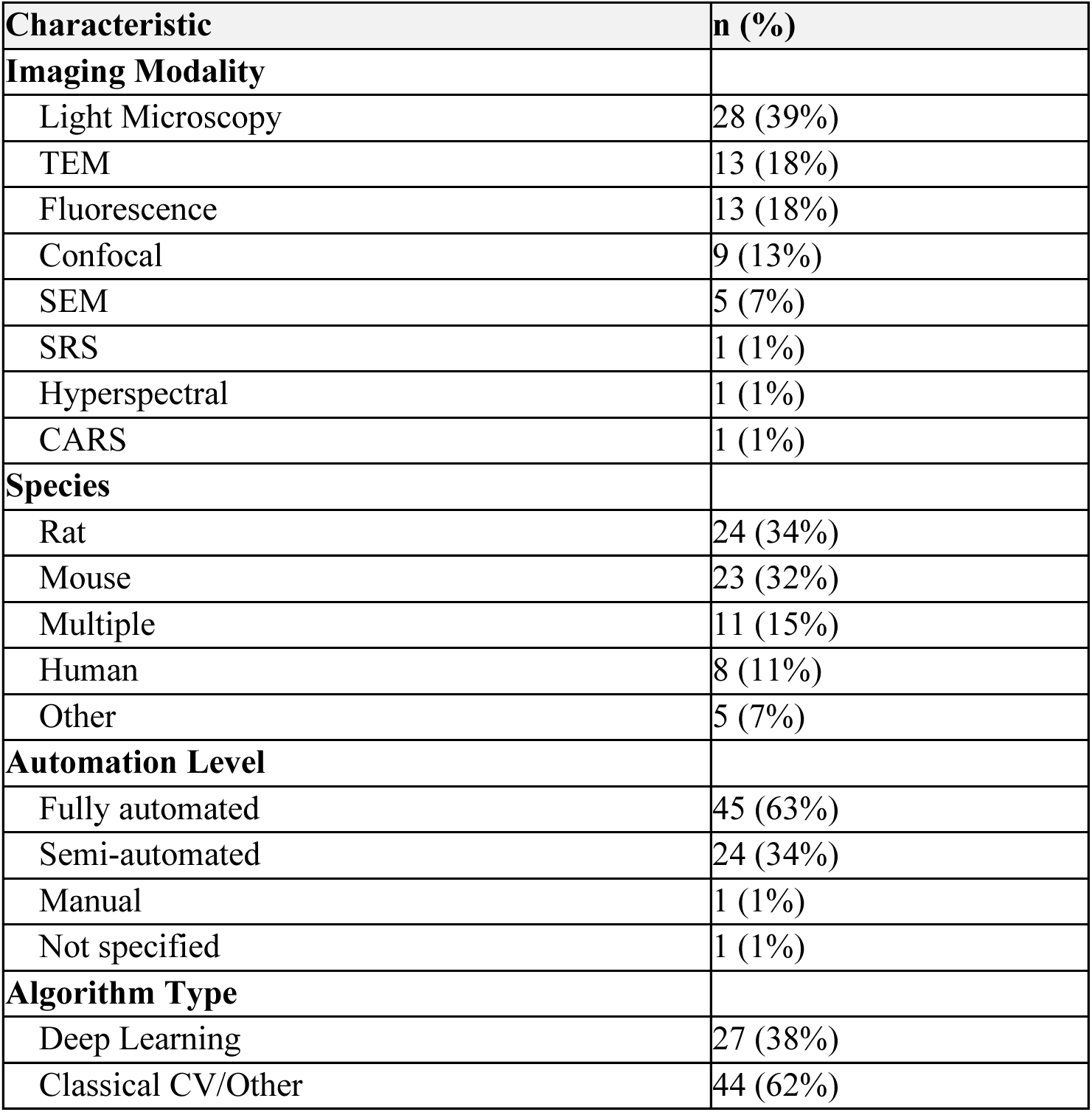
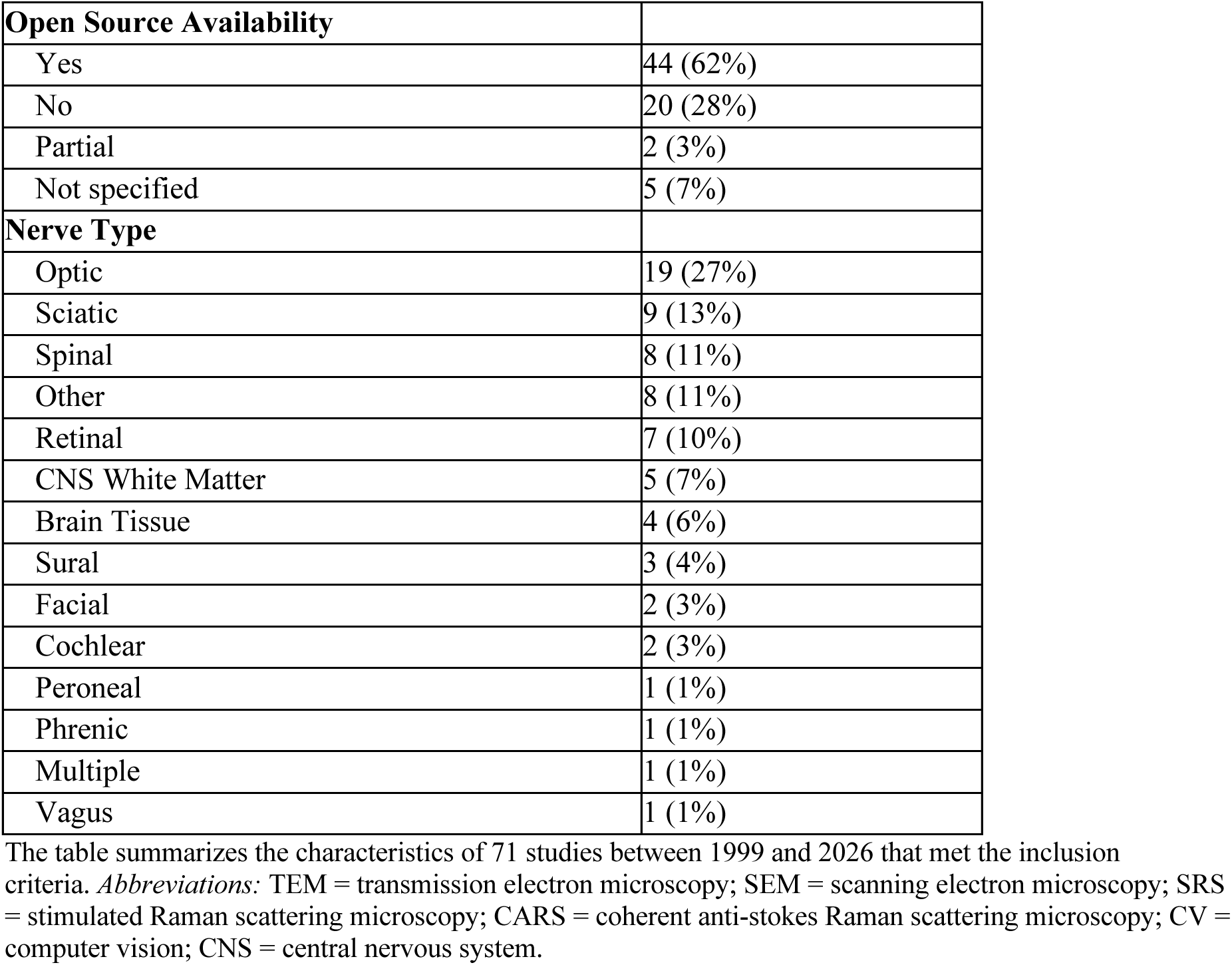
Characteristics of Included Studies.

Scanning electron microscopy (SEM) accounted for 7% (5/71), with the remainder comprising stimulated Raman spectroscopy, coherent anti-Stokes Raman scattering, and hyperspectral imaging (1 study each).

Rat was the most frequently studied species (34%, 24/71), followed by mouse (32%, 23/71), multiple species (15%, 11/71), and human (11%, 8/71). The majority of tools were reported as fully automated (63%, 45/71), with 34% (24/71) described as semi-automated and one study each using manual or unspecified methods.

Optic nerve was the most common tissue target (27%, 19/71), followed by sciatic nerve (13%, 9/71) and retinal tissue (10%, 7/71). Other nerve types included spinal cord (11%, 8/71), CNS white matter (7%, 5/71), brain tissue (6%, 4/71), sural nerve (4%, 3/71), and facial nerve (3%, 2/71).

#### Publication Trends

Publication of nerve morphometry tools accelerated markedly after 2017, coinciding with the adoption of deep learning (**Figure 2**). Before 2017, all published tools used classical CV or other non-deep-learning approaches. Deep learning tools for nerve morphometry first appeared in 2017 and represented over 55% of publications from 2020 onward. Of the 71 total studies, 27 (38%) used deep learning and 44 (62%) used classical CV or other methods.

**Figure 2.**
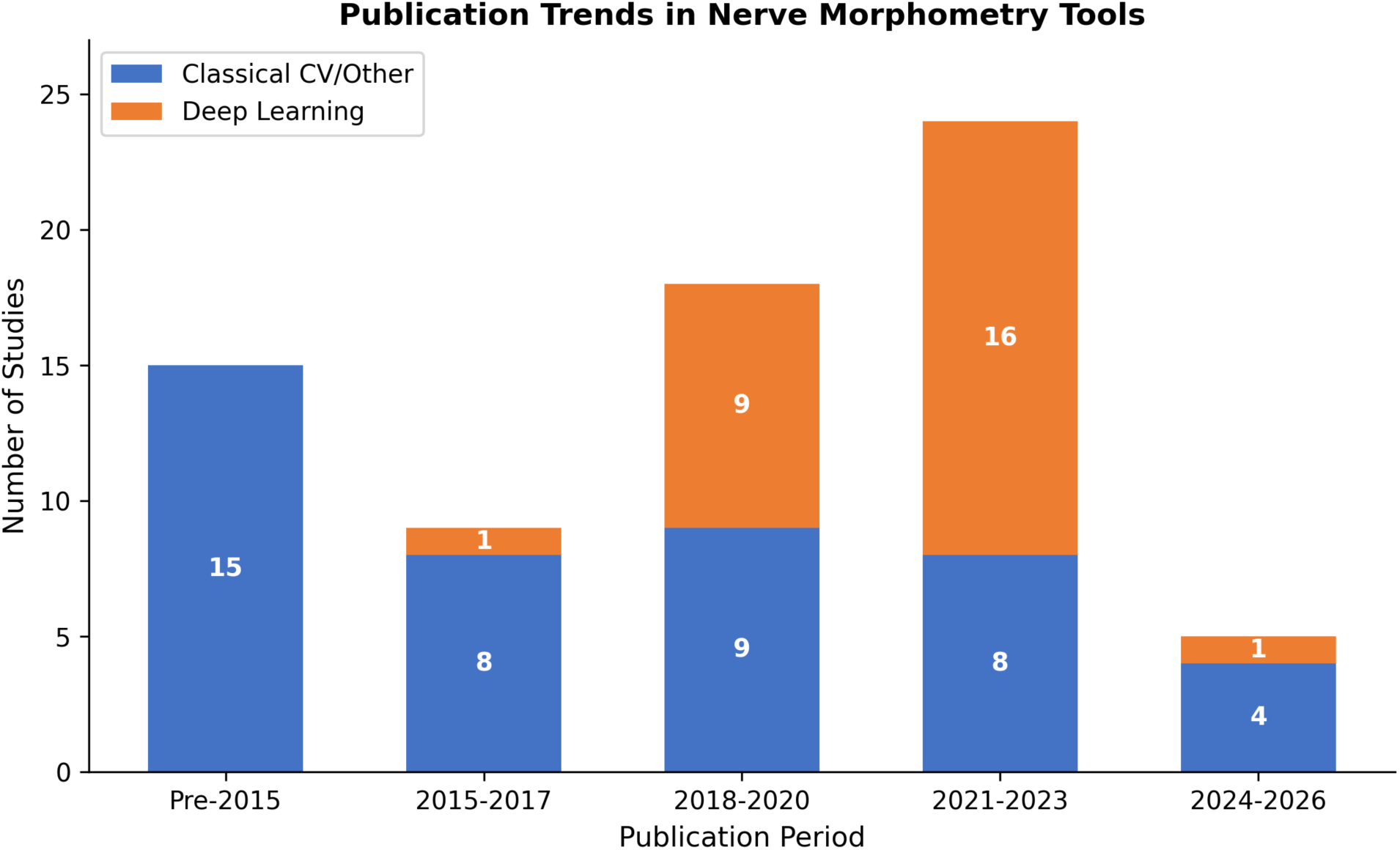
Publication Trends in Nerve Morphometry Tools. Stacked bar chart showing the number of published nerve morphometry tools by publication period. n = 71 studies spanning 1999 to 2026. *Abbreviation:* CV = computer vision.

#### Tool Capabilities

Axon counting was the most commonly provided output (73%, 52/71), followed by size distribution analysis (58%, 41/71), axon diameter measurement (48%, 34/71), and axon area quantification (48%, 34/71) (**Table 2**, **Figure 3**). Less frequently reported outputs included g-ratio (35%, 25/71), fiber diameter (35%, 25/71), fiber count (32%, 23/71), myelin thickness (31%, 22/71), fiber density (30%, 21/71), and myelin area (27%, 19/71).

**Figure 3.**
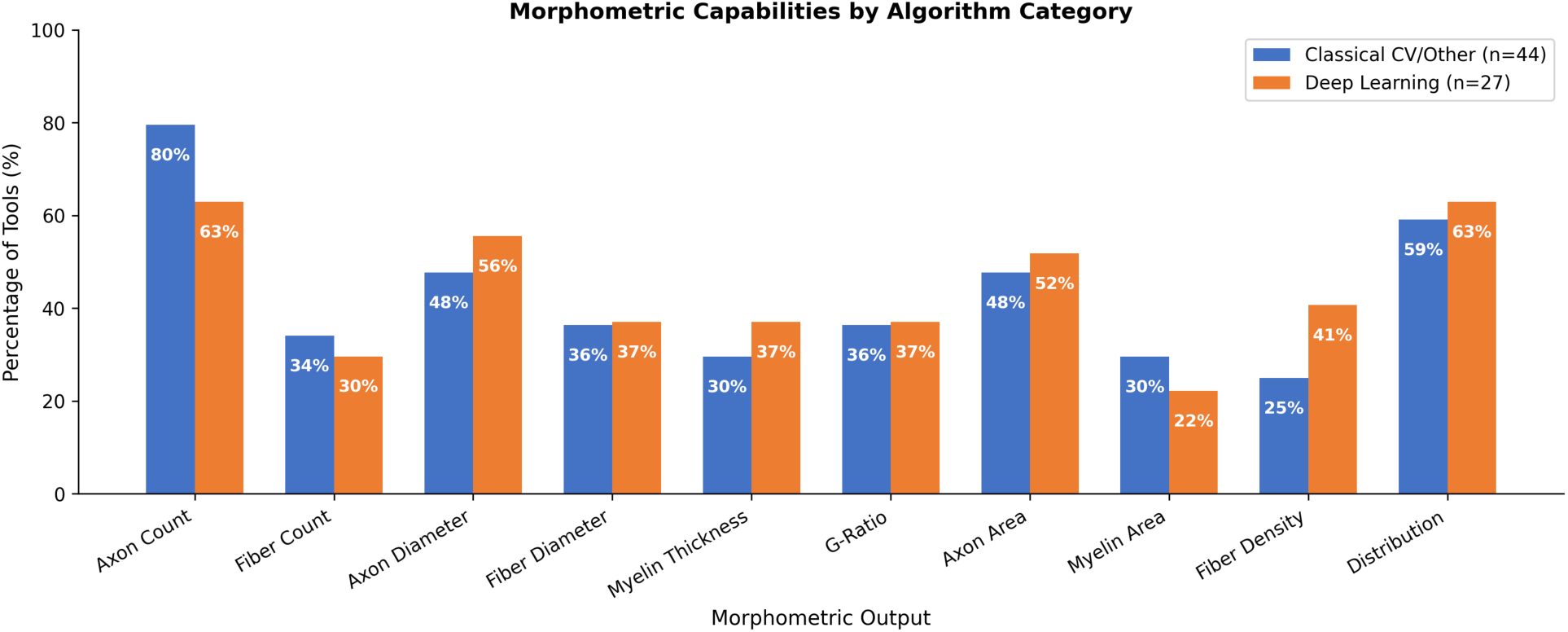
Morphometric Capabilities by Algorithm Category. Grouped bar chart compares the percentage of tools offering each morphometric output. *Abbreviations:* CV = computer vision.

**Table 2.**
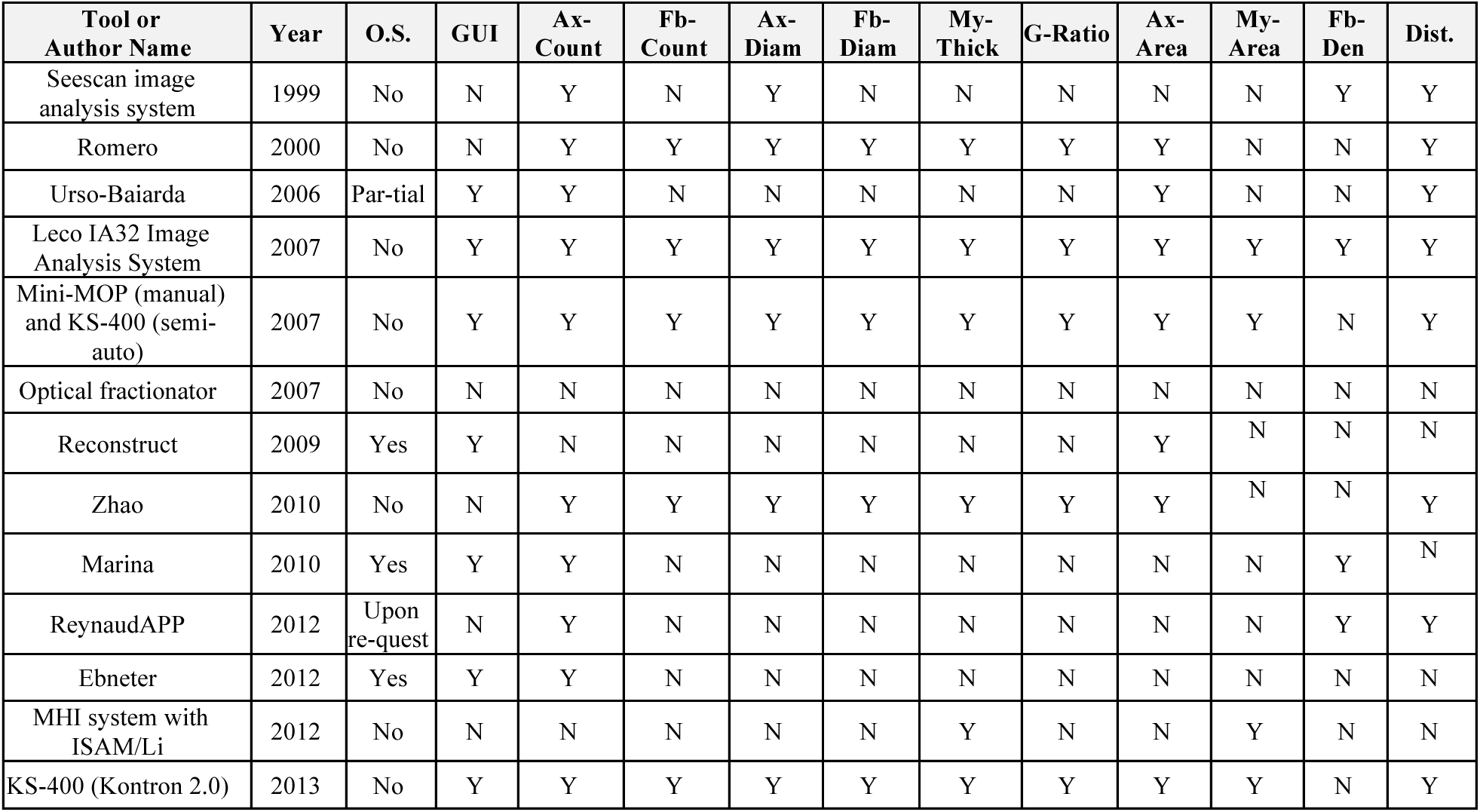
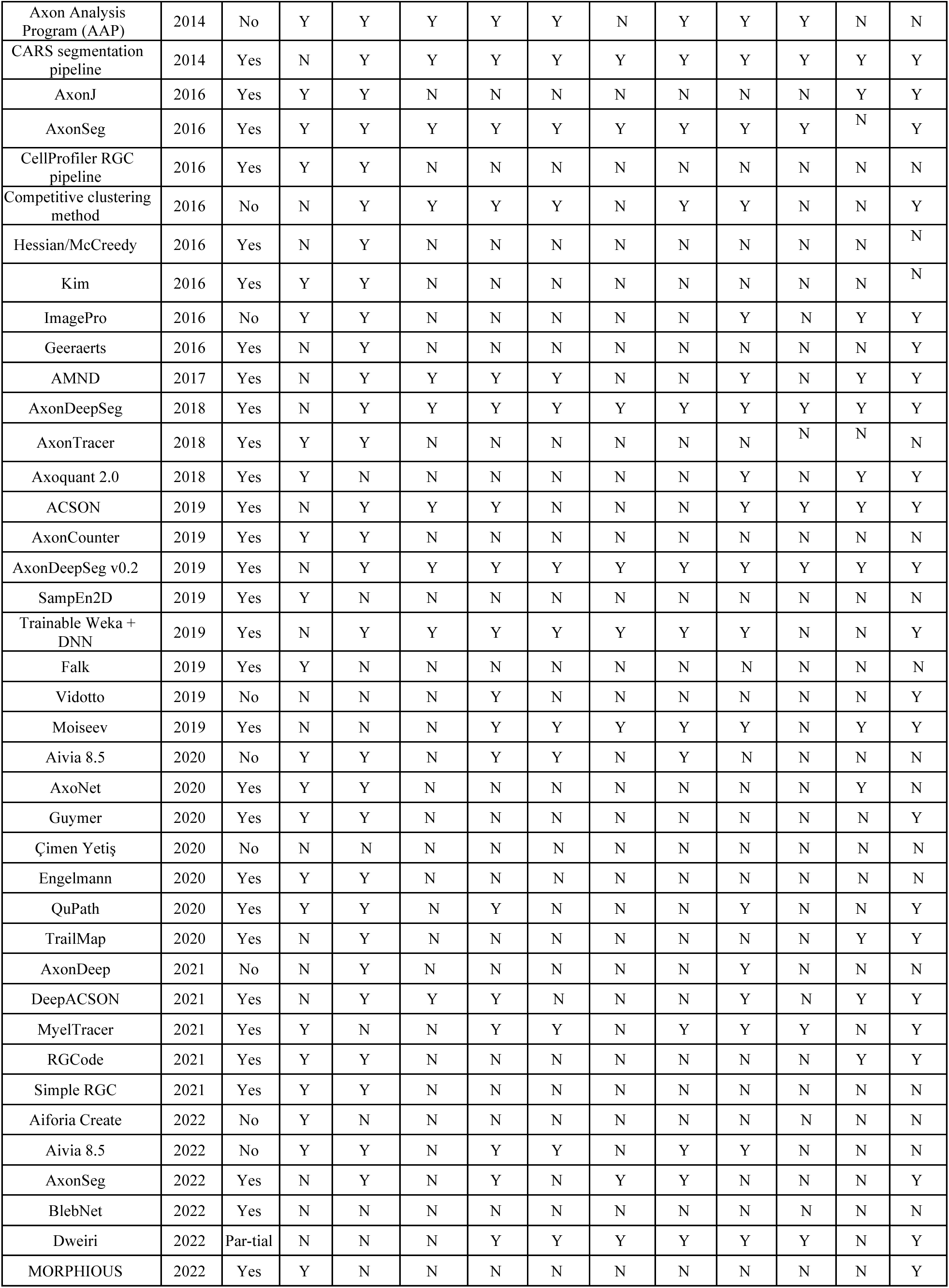
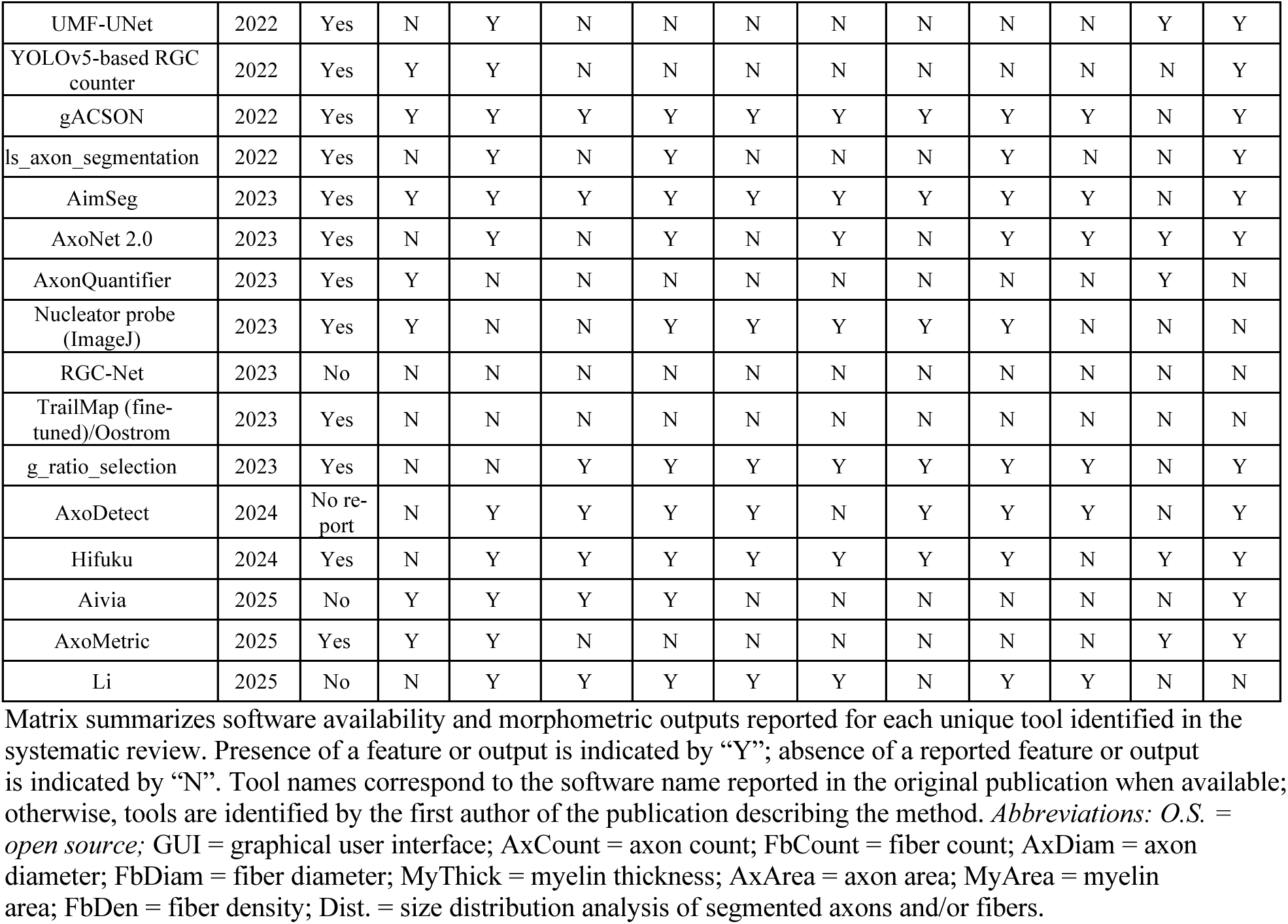
Tool Capabilities.

Classical CV tools were more likely than deep learning tools to provide axon counting (80% vs. 63%, respectively) but less likely to measure axon diameter (48% CV vs. 56% DL) or fiber density (25% CV vs. 41% DL). G-ratio and fiber diameter measurements were comparable between categories (36% DL vs. 37% CV for both).

Open-source code was available for 62% (44/71) of tools, with an additional 3% (2/71) providing partial source code. A GUI was available for 54% (38/71) of tools. Among the 27 deep learning tools, 74% (20/27) were open source, compared to 52% (23/44) of classical CV tools.

### Independent Benchmarking

#### Overall Performance

Performance metrics for all 18 benchmarked tools across both datasets are presented in **Table 3** and **Figure 4**. Average MAPE across tools ranged from 32.9% (Marina, CV) to 233.2% (AxonJ, CV). The top five tools by average MAPE were: Marina (32.9%), ADS Generalist (41.4%), AxoNet 2.0 (43.8%), AxoNet (44.3%), and ADS Bright-Field Light (44.8%). These top performing tools consisted of four DL tools and one CV tool.

**Figure 4.**
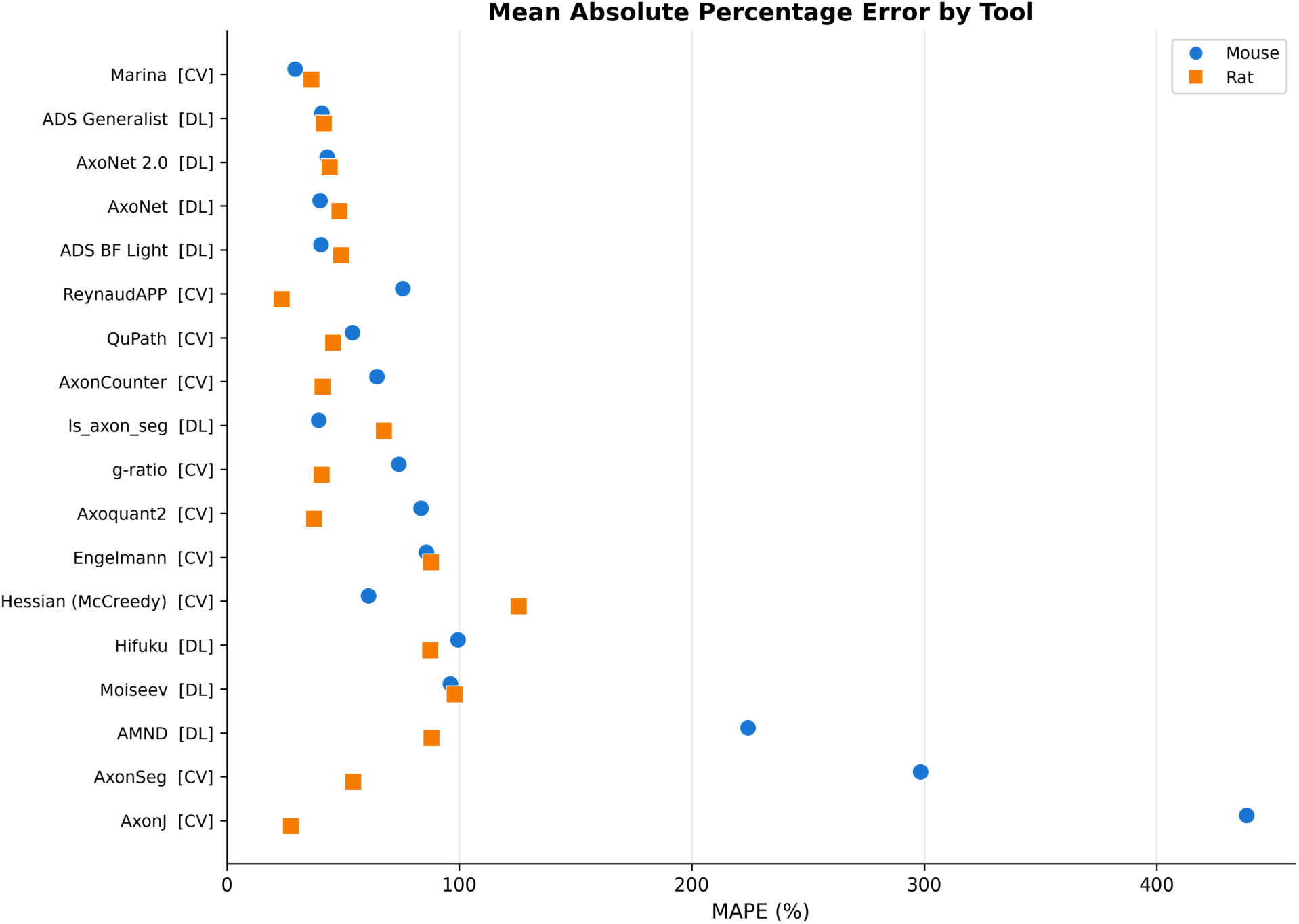
Mean Absolute Percentage Error (MAPE) by Tool. MAPE for 18 benchmarked tools across paraphenylenediamine (PPD)-stained mouse and rat datasets is shown on a forest plot. *Abbreviations:* CV = computer vision; DL = deep learning.

**Table 3.** Benchmarking Results.

| Tools |  |  | Mouse |  |  |  |  |  |  | Rat |  |  |  |  |  |  | Avg MAPE |
| --- | --- | --- | --- | --- | --- | --- | --- | --- | --- | --- | --- | --- | --- | --- | --- | --- | --- |
| Rk | Name (year) | Ct | MAPE | r | MAE | RMSE | Mdn ratio | R<sup>2</sup> | N<sub>mt</sub> | MAPE | r | MAE | RMSE | Mdn ratio | R<sup>2</sup> | N<sub>mt</sub> |  |
| 1 | Marina (2010) | CV | 29.3 | 0.73 | 5405.0 | 6736.2 | 1.07 | 0.01 | 75 | 36.4 | 0.82 | 57.1 | 73.6 | 0.70 | 0.31 | 44 | 32.85 |
| 2 | ADS Generalist (2018) | DL | 40.9 | 0.54 | 8239.9 | 10011.2 | 0.69 | -1.20 | 75 | 41.8 | 0.75 | 60.1 | 76.4 | 0.63 | 0.25 | 44 | 41.35 |
| 3 | AxoNet 2.0 (2023) | DL | 43.2 | 0.56 | 8115.1 | 9590.9 | 1.01 | -1.0 | 75 | 44.3 | 0.91 | 63.2 | 74.9 | 0.58 | 0.28 | 44 | 43.75 |
| 4 | AxoNet (2020) | DL | 40.1 | 0.70 | 7609.6 | 9529.7 | 1.28 | -0.99 | 75 | 48.5 | 0.92 | 73.8 | 87.3 | 0.51 | 0.02 | 44 | 44.3 |
| 5 | ADS BF<br>Light<br>(2018) | DL | 40.4 | 0.51 | 7620.6 | 9338.2 | 0.72 | -0.89 | 74 | 49.2 | 0.90 | 74.9 | 90.3 | 0.49 | -0.05 | 44 | 44.8 |
| 6 | Reynaud-APP<br>(2012) | CV | 75.7 | 0.39 | 14368.5 | 18554.2 | 1.39 | -6.55 | 75 | 23.6 | 0.92 | 30.3 | 37.2 | 1.12 | 0.82 | 44 | 49.65 |
| 7 | QuPath<br>(2020) | CV | 54.0 | 0.53 | 9989.3 | 13413.9 | 1.17 | -2.94 | 75 | 45.7 | 0.59 | 83.2 | 110.9 | 0.49 | -0.58 | 44 | 49.85 |
| 8 | Axon-Counter<br>(2017) | CV | 64.5 | 0.58 | 11577.3 | 13323.7 | 1.4 | -2.89 | 75 | 41.2 | 0.90 | 46.3 | 57.4 | 1.21 | 0.58 | 44 | 52.85 |
| 9 | ls_axon_seg<br>(2022) | DL | 39.5 | 0.49 | 7726.0 | 9503.2 | 0.80 | -0.95 | 74 | 67.6 | 0.23 | 108.4 | 139.9 | 0.17 | -1.51 | 44 | 53.55 |
| 10 | g-ratio<br>(2023) | CV | 74.0 | 0.15 | 14713.0 | 17138.7 | 0.38 | -5.44 | 75 | 40.8 | 0.69 | 69.1 | 92.7 | 0.60 | -0.10 | 44 | 57.4 |
| 11 | Axoquant2<br>(2018) | CV | 83.5 | 0.67 | 14702.1 | 15639.5 | 0.12 | -4.20 | 19 | 37.6 | 0.64 | 74.6 | 102.7 | 0.61 | -0.35 | 44 | 60.55 |
| 12 | Engelmann<br>(2020) | CV | 85.8 | 0.10 | 18609.6 | 19827.4 | 0.11 | -7.62 | 75 | 87.9 | 0.48 | 136.0 | 156.9 | 0.05 | -2.16 | 44 | 86.85 |
| 13 | Hessian-McCreedy<br>(2016) | CV | 60.9 | 0.47 | 10892.9 | 14208.4 | 1.29 | -3.42 | 75 | 125.5 | 0.53 | 123.6 | 144.3 | 1.88 | -1.67 | 44 | 93.2 |
| 14 | Hifuku<br>(2024) | DL | 99.3 | 0.14 | 21325.3 | 22364.6 | 0.004 | -9.96 | 75 | 87.5 | 0.81 | 135.8 | 155.5 | 0.12 | -2.10 | 44 | 93.4 |
| 15 | Moiseev<br>(2019) | DL | 96.2 | 0.26 | 20705.8 | 21740.2 | 0.03 | -9.36 | 75 | 98.0 | -0.22 | 155.8 | 179.3 | 0.01 | -3.12 | 44 | 97.1 |
| 16 | AMND (2021) | DL | 224.2 | 0.28 | 42182.6 | 48646.9 | 2.82 | -50.87 | 75 | 88.0 | 0.55 | 140.7 | 162.9 | 0.09 | -2.40 | 44 | 156.1 |
| 17 | AxonSeg (2017) | CV | 298.4 | 0.35 | 57949.6 | 66960.6 | 3.54 | -97.27 | 75 | 54.4 | 0.83 | 90.6 | 111.7 | 0.43 | -0.6 | 44 | 176.4 |
| 18 | AxonJ (2016) | CV | 438.6 | -0.12 | 72733.9 | 174634.3 | 1.66 | -667.39 | 75 | 27.7 | 0.84 | 38.5 | 52.3 | 0.91 | 0.65 | 44 | 233.2 |

Performance metrics for 18 tools across paraphenylenediamine (PPD)-stained mouse and rat datasets. Tools were ranked by average mean absolute percentage error (MAPE). *Abbreviations:* Rk = rank; Ct = category; r = Pearson r; MAE = mean absolute error; RMSE = root mean square error; mdn ratio = median ratio; N\_mt = N\_matched; Avg MAPE = average mean absolute percentage error.

Pearson correlation between predicted and ground truth counts varied widely (**Figure 5**). On the mouse dataset, Marina achieved the highest correlation (r = 0.73), followed by AxoNet (r = 0.70) and Axoquant 2.0 (r = 0.67). On the rat dataset, AxoNet and ReynaudAPP both achieved the highest correlation (r = 0.92), followed by AxoNet 2.0 (r = 0.91) and ADS Bright-Field Light (r = 0.90).

**Figure 5.**
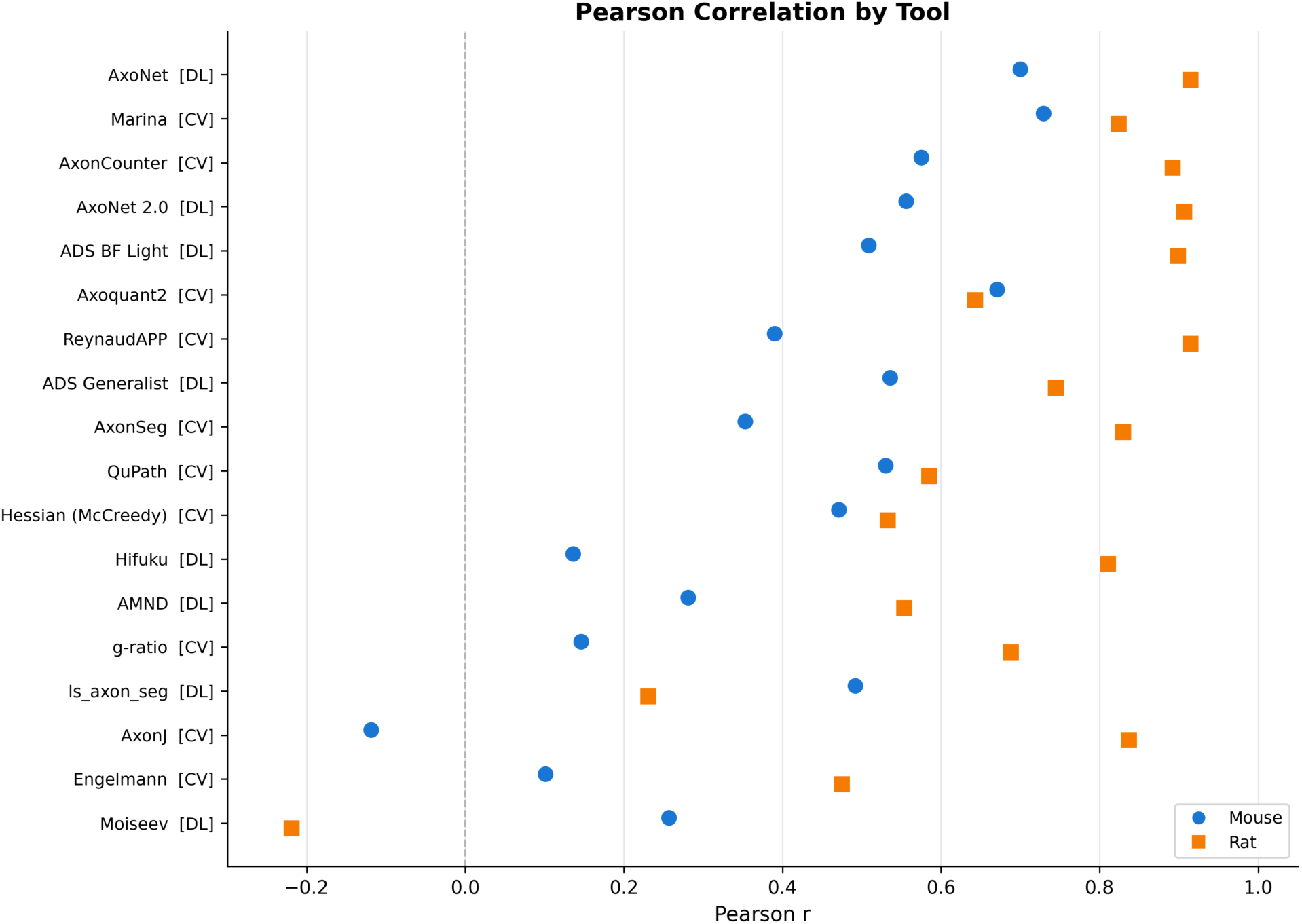
Pearson Correlation (r) by tool. The forest plot of Pearson correlation r for 18 tools shows varied results between predicted and ground truth counts using paraphenylenediamine (PPD)-stained mouse and rat optic nerve image datasets. *Abbreviations:* CV = computer vision; DL = deep learning.

#### Dataset-Specific Performance

Performance varied substantially between datasets (**Figure 6, Supplementary Figure S1**). On the mouse dataset, Marina achieved the lowest MAPE (29.3%), followed by ls_axon_segmentation (39.5%), AxoNet (40.1%), ADS Bright-Field Light (40.4%), and ADS Generalist (40.9%). On the rat dataset, ReynaudAPP achieved the lowest MAPE (23.6%) followed by AxonJ (27.7%), Marina (36.4%), Axoquant 2.0 (37.6%), and g-ratio (40.8%).

**Figure 6.**
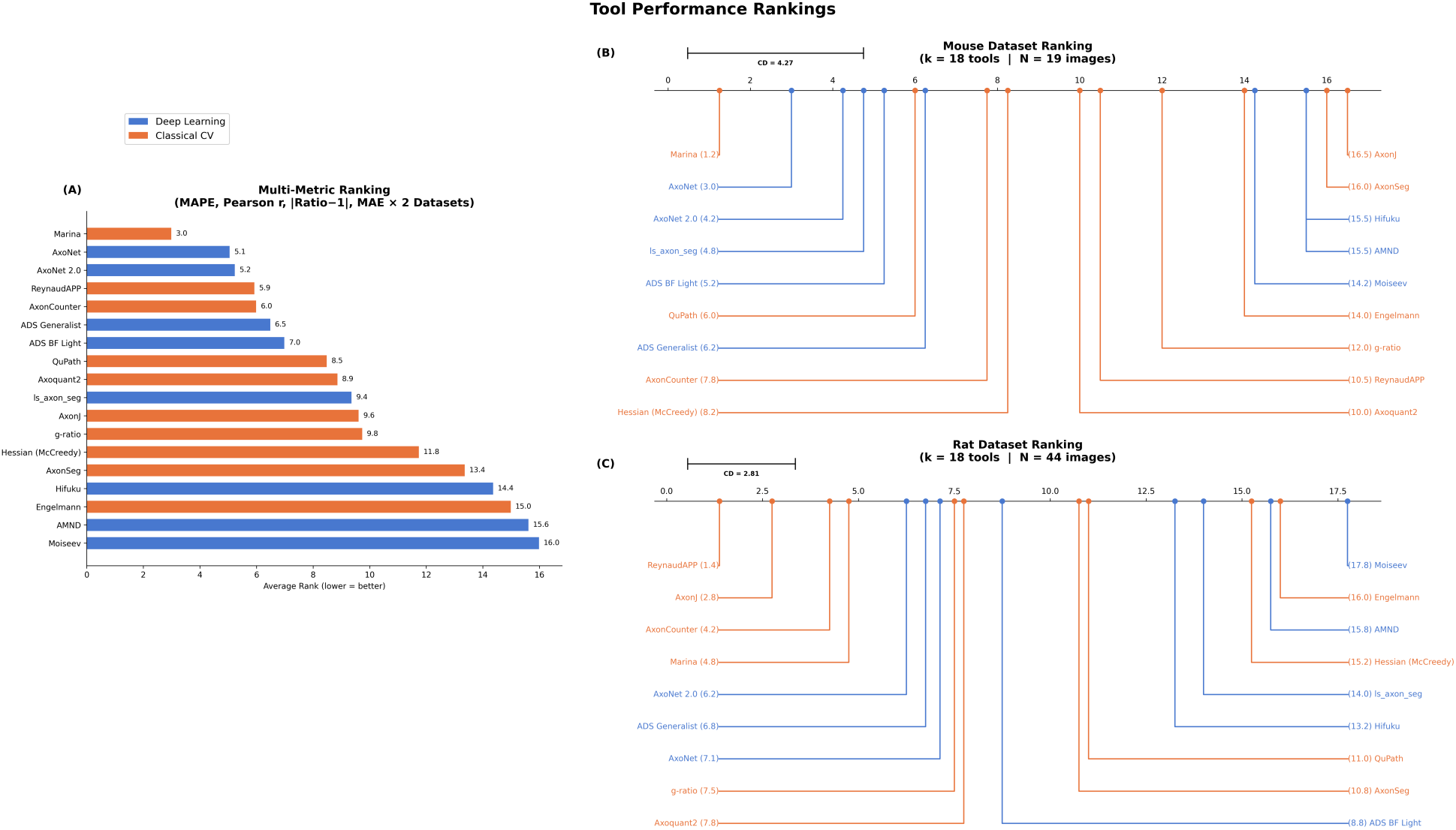
Friedman-Nemenyi critical difference (CD) diagrams comparing tool rankings. (A) Multi-metric rankings across both datasets generated by combining performance metrics of mean absolute percentage error (MAPE), Pearson correlation, absolute prediction bias (|Ratio – 1|), and mean absolute error (MAE). (B) Rankings based on performance in the mouse dataset. (C) Rankings based on performance in the rat dataset. Lower average rank indicates better performance. The CD represents the minimum separation in average rank required for statistical significance according to the Nemenyi post-hoc test. Tools within the CD range do not differ significantly in performance (p > 0.05), whereas tools separated by more than the CD differ significantly (p <0.05).

Several tools showed divergent performance across the two datasets. AxonJ ranked last on the mouse dataset (MAPE 438.6%) but second on the rat dataset (MAPE 27.7%). Conversely, ls_axon_segmentation ranked second on the mouse dataset (MAPE 39.5%) but 13th on rat dataset (MAPE 67.6%). These disparities suggest sensitivity to species, sample quality, or image characteristics (**Supplementary Figure S2**).

#### Bias Assessment

The median predicted-to-ground-truth ratio revealed systematic biases (**Supplementary Figure S3**). On the mouse dataset, ten tools showed a tendency to overcount (ratio > 1.0: 3.54 AxonSeg - 1.01 AxoNet 2.0), while eight tools undercounted (ratio < 1.0: 0.004 Hifuku – 0.80 ls_axon_seg). On the rat dataset, three tools Hessian/McCreedy consistently overcounted (Hessian/McCreedy 1.88, AxonCounter 1.21, ReynaudAPP 1.12), and all other tools had ratios below 1.0 (0.01 Moiseev – 0.91 AxonJ), indicating a general tendency toward underestimation on rat tissue.

#### Rank-Based Statistical Comparison

Friedman test results indicated statistically significant differences in tool rankings across images (p < 0.001 for both datasets individually and combined). Nemenyi post-hoc analysis identified clusters of tools with statistically indistinguishable performance (**Figure 6**). In the combined analysis, the pairwise comparison within the top performing tools (Marina, AxoNet, AxoNet 2.0, ReynaudAPP, and AxonCounter) reached no significance at p < 0.05.

On the mouse dataset alone, the top cluster included Marina, AxoNet, Axonet 2.0, ls_axon_segmentation, and ADS Bright-Field Light. On the rat dataset, the top cluster included ReynaudAPP, AxonJ, AxonCounter, Marina, and Axonet 2.0. Only Marina and AxoNet 2.0 appeared in the top statistical cluster for both datasets.

#### Segmentation Performance and Qualitative Comparison

Segmentation performance was evaluated for tools that produced pixel-level output masks (**Table 4**). Performance varied across models and datasets. AMND achieved a Dice coefficient of 0.40 on the rat dataset with high recall (0.91) but low precision (0.28), indicating over segmentation of background regions while still capturing most true axons. Moiseev demonstrated near-complete failure to recognize optic nerve axons on the rat dataset (Dice = 0.0007).

**Table 4.** Segmentation Performance and Generalizability Gap.

| Tools |  |  | Published Values |  |  |  |  | Independent Mouse Dataset |  | Independent Rat Dataset |  |  |  |
| --- | --- | --- | --- | --- | --- | --- | --- | --- | --- | --- | --- | --- | --- |
| Name | Cat | Rk | r | r Source | Dice | Dice Source | Species/Nerve | r | r Diff | r | Dice | r Diff | Dice Diff |
| Marina | CV | 1 | 0.98 | Marina 2010 | NR | NR | Rat/Optic | 0.73 | 0.24 | 0.82 | - | 0.14 | - |
| ADS Generalist | DL | 2 | NR | NR | 0.91 | Zaimi 2018 | Rat/Human/Mouse/Spinal cord | 0.54 | - | 0.75 | 0.43 | - | 0.48 |
| AxoNet 2.0 | DL | 3 | 0.92 | Goyal 2023 | 0.81 | Goyal 2023 | Rat/Mouse/NHP/Optic | 0.56 | 0.36 | 0.91 | 0.34 | 0.01 | 0.47 |
| AxoNet | DL | 4 | 0.97 | Ritch 2020 | NR | NR | Rat/NHP/Optic | 0.7 | 0.27 | 0.92 | - | 0.05 | - |
| ADS BF Light | DL | 5 | NR | NR | 0.91 | Zaimi 2018 | Rat/Human/Mouse/Spinal cord | 0.51 | - | 0.90 | 0.44 | - | 0.47 |
| Reynaud-APP | CV | 6 | 0.98 | Reynaud 2012 | NR | NR | NHP/Optic | 0.39 | 0.59 | 0.92 | - | 0.06 | - |
| QuPath | CV | 7 | 0.96 | Mysona 2020 | NR | NR | Rat/Optic | 0.53 | 0.43 | 0.59 | - | 0.37 | - |
| Axon-Counter | CV | 8 | NR | NR | NR | NR | Mouse/Optic | 0.58 | - | 0.89 | - | - | - |
| ls_axon_seg | DL | 9 | NR | NR | 0.77 | Mordhorst 2022 | Human/Corpus callosum | 0.49 | - | 0.23 | 0.15 | - | 0.62 |
| g-ratio | CV | 10 | NR | NR | 0.86 | Thomson 2023 | Mouse/Sciatic | 0.15 | - | 0.69 | - | - | - |
| Axoquant2 | CV | 11 | NR | NR | NR | NR | Mouse/DRG | 0.67 | - | 0.64 | - | - | - |
| Engelmann | CV | 12 | 0.94 | Engelmann 2020 | NR | NR | Human/Facial | 0.10 | 0.84 | 0.48 | - | 0.47 | - |
| Hessian (McCreedy) | CV | 13 | 0.85 | Mc-Creedy 2016 | NR | NR | Mouse/Spinal cord | 0.47 | 0.38 | 0.53 | - | 0.32 | - |
| Hifuku | DL | 14 | NR | NR | NR | NR | Human/Sural | 0.14 | - | 0.81 | - | - | - |
| Moiseev | DL | 15 | NR | NR | 0.83 | Moiseev 2019 | Mouse/Sciatic | 0.26 | - | -0.22 | 0.001 | - | 0.83 |
| AMND | DL | 16 | NR | NR | NR | NR | Human/Sural | 0.28 | - | 0.55 | 0.41 | - | - |
| AxonSeg | CV | 17 | NR | NR | NR | NR | Rat/Cat/Spinal cord | 0.35 | - | 0.83 | - | - | - |
| AxonJ | CV | 18 | 0.95 | Zarei 2016 | NR | NR | Mouse/Optic | -0.12 | 1.07 | 0.84 | - | 0.11 | - |

Pearson correlation coefficients (r) for axon counting and Dice coefficients for segmentation obtained in the present benchmarking study were compared with values reported in the original publications for 18 evaluated tools. Differences were calculated as benchmark performance minus published performance. “NR” indicates that the metric was not reported in the original publication. “-” indicates that a comparison could not be performed because no corresponding published reference value or independent benchmarking value was available. *Abbreviations:* Cat = category; Rk = rank; r = Pearson correlation (r); Diff = difference; NHP = nonhuman primate.

Similarly, ls_axon_seg showed reduced segmentation performance on the rat dataset (Dice = 0.17), despite stronger counting performance on the mouse model.

Visual comparison of representative segmentation outputs is shown in **Figure 7**. ADS, AxonSeg, and ls_axon_seg generated pixel-level axon and myelin masks, whereas AxoNet 2.0 produced pixel-level axon probability maps that were converted to binary axon masks for analysis. These tools captured the overall axon distribution pattern, though differences were visible in regions of densely packed or damaged axons. Segmentation masks identify individual axon and myelin profiles, in contrast to tools like Marina and ReynaudAPP that output aggregate count estimates without spatial localization, precluding pixel-level comparison. Among these, Marina achieved the lowest MAPE value in the benchmark (**Table 3**), demonstrating that accurate axon quantification does not require full segmentation.

**Figure 7.**
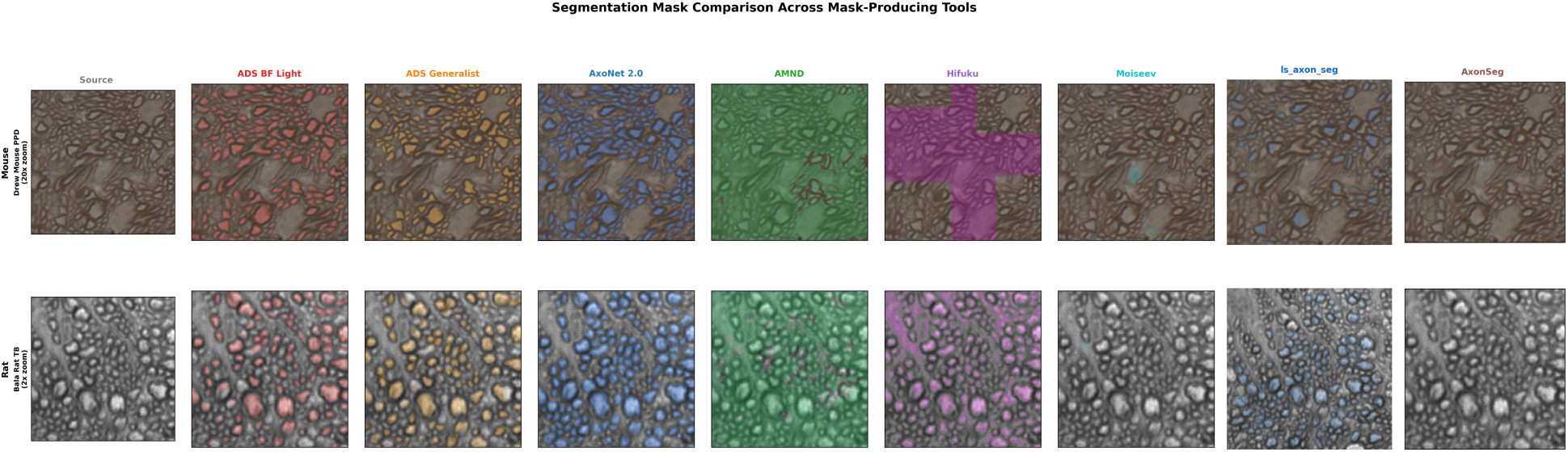
Segmentation mask comparison across mask-producing tools. Predicted segmentation masks from eight mask-producing tools overlaid on representative source images from paraphenylenediamine (PPD)-stained mouse (top row, 20x zoom of 550x550 pixel crop from center of image 031109_39L) and rat (bottom row, full 256x256 pixel patch at native resolution) datasets. Tools shown are ADS Bright-Field Light (red), ADS Generalist (orange), AxoNet 2.0 (blue), AMND (green), Hifuku (purple), Moiseev (teal), ls_axon_seg (dark blue), and AxonSeg (brown). Masks are displayed as colored overlays (alpha = 0.25) on the original grayscale source images. The figure illustrates qualitative differences in spatial localization across tools.

#### Generalizability Analysis

To assess generalizability, performance on the independent benchmarking datasets was compared with the validation metrics reported in the original publications (**Table 4, Supplementary Figure S4**). Seven tools reported Pearson correlation and six tools reported Dice coefficient for axon counts compared to their ground truth data. Across all evaluated tools, performance on the independent mouse and rat optic nerve datasets was consistently lower than reported by the original authors for both Pearson r and Dice coefficient.

## Discussion

This study presents the first systematic review of automated nerve morphometry tools alongside an independent benchmarking evaluation on shared optic nerve datasets. Across 71 studies, we observed a rapid growth in the field from 1999 to 2026, particularly following the adoption of deep learning methods since 2017.^21^ Despite these advances, independent benchmarking revealed substantial performance variability across datasets and consistently lower performance than originally reported, highlighting the continued challenge of model generalizability.

Contrary to expectations,^32,33^ deep learning methods did not uniformly outperform classical CV methods. Several classical methods achieved performance comparable to top-performing deep learning models, suggesting that algorithm class alone is not the primary determinant of accuracy. These findings suggest that well-tuned classical algorithms remain competitive for axon counting, particularly when the target tissue is well-stained and relatively uniform, such as seen in optic nerve histology.^5,17,27^ That said, the top five tools from independent benchmarking included four DL methods (ADS Generalist, AxoNet 2.0, AxoNet, ADS Bright-Field Light).

Rather than replacing classical methods, deep learning appears to have expanded the range of viable tools available, particularly for applications requiring adaptability across imaging conditions.^15,34^

The most consistent finding across the benchmark was the limited generalizability. Performance varied between mouse and rat datasets despite both datasets using optic nerve histology, and all evaluated tools performed worse on independent data than reported in their original publications when comparable metrics were available. Such a challenge in biomedical image analysis has been reported in other studies, in which models often experience worse performance when applied to data acquired under different protocols, species, or imaging systems.^35–38^

In our study, several of the tools trained on peripheral nerve tissue demonstrated limited transferability to optic nerve images, particularly AMND, Hifuku, and Moiseev. AMND was trained on sciatic and sural nerve tissue,^21^ Hifuku trained for sural nerve fascicle-level segmentation,^19^ and Moiseev trained for sciatic nerve myelin segmentation.^20^ These results demonstrate that tools trained and validated on peripheral nerve tissue do not reliably transfer to optic nerve without retraining, even when the underlying architectures are capable. Interestingly, models developed using optic nerve tissue, such as AxoNet, have reported successful transfer across species without retraining, including application to mouse and non-human primate optic nerve images.^13^ Likewise, the AxonDeepSeg framework trained on heterogeneous datasets spanning multiple tissue preparations and species showed relatively strong performance across our benchmark datasets and in other studies^15,39,40^ Together, these findings emphasize the importance of external validation and suggest that future development should prioritize training with robust sources rather than optimizing performance using a single dataset.

Tool selection should also consider the desired biological outputs rather than axon-counting performance alone. While axon count was the most commonly reported metric, fewer than 40% of tools reported myelin-related measurements, and only a minority provided comprehensive morphometric analysis **(Supplementary Figure S5**). Metrics such as axon diameter distributions, myelin thickness, fiber density, and g-ratio can provide complementary information regarding axonal health, myelination status, and ultimately disease progression.^4,41–43^ Because axon count alone does not fully characterize optic nerve health, access to these additional morphometric measurements may be an important consideration when selecting a tool. Among the benchmarked tools, AxonDeepSeg and AxonSeg were notable for providing both axon and myelin segmentation masks with downstream morphometric measurements.^9,17^ While ls_axon_seg also generated both axon and myelin segmentations, it was primarily developed for axon radius estimation.^18^

The review also revealed heterogeneity in validation practices. Studies varied widely dataset size, annotation procedures, outcome metrics, and reporting practices (**Supplementary Table S3**). Many studies relied on internal validation datasets, and few reported external validation or interobserver agreements. These differences limit direct comparison across tools and emphasize the need for standardized benchmark datasets and reporting guidelines for further tool development. Nevertheless, both CV and DL approaches substantially reduce analysis time compared with manual annotation, with reported reductions ranging from approximately 40- 80%.^20,44–46^ Despite variability in performance across tools, automated morphometry remains essential for enabling large-scale quantitative analyses as publicly available imaging datasets and shared data repositories continue to expand.

Several limitations should be acknowledged. First, the benchmarking was limited to axon counting accuracy because this was the only output common to all 18 tools; tools that also produce segmentation masks or morphometric measurements were not compared on those outputs. Second, ground truth was based on manual counts rather than pixel-level annotations, which prevents evaluation of segmentation quality at the individual axon level. Third, the two datasets represent only PPD-stained mouse and rat optic nerve; generalization to other species (e.g., primate, human), staining protocols (e.g., toluidine blue, osmium, immunohistochemistry), or nerve types (e.g., sciatic, sural) were not tested. Fourth, comparison of our benchmarking results with originally published performance metrics was possible for only 13 of 18 tools. 8 reported count correlation, 6 reported Dice coefficient, with some overlap between groups (**Table 4 and Supplementary Figure S4**). The remaining 5 tools had no published quantitative metrics suitable for direct comparison. Furthermore, original publications generally reported aggregate statistics rather than per-image data, limiting comparisons to summary-level concordance and precluding paired analysis. Lastly, tool performance may depend on parameter tuning, preprocessing choices, and threshold selection.^14,34,47–49^ While we used default or recommended settings for each tool for reproducibility, additional optimization on each dataset could improve performance for some methods.

In summary, automated neural tissue morphometry has matured considerably from early threshold-based image analysis to deep learning frameworks, yet external validation remains limited. Our benchmarking demonstrates that both classical and deep learning approaches can achieve strong performance for axon quantification, but that accuracy depends strongly on dataset characteristics and similarity to the original training data. Therefore, tool selection should be guided by target tissue characteristics and the specific morphometric outputs required for a given study rather than algorithm class alone. These findings highlight the need for standardized benchmark datasets, broader external validation, and development of models for cross-species and protocol generalization.

## Meeting Presentation

This material is under consideration for presentation at the American Academy of Ophthalmology Annual Meeting, 2027.

## Abbreviations

CV: computer vision
DL: deep learning
GUI: graphical user interface
PRISMA: preferred reporting items for systematic reviews and meta-analysis
CNN: convolutional neural network
ADS: AxonDeepSeg
PPD: paraphenylenediamine
MAPE: mean absolute percentage error
MAE: mean absolute error
RMSE: root mean square error.

## Declaration of Generative AI and AI-assisted technologies in the writing process

During the preparation of this work the author(s) used publicly available original repositories of semi-automated and automated axon and myelin counting tools listed in **Supplementary Table S3** for independent benchmarking. After using this tool/service, the authors reviewed and edited the content as needed and take full responsibility for the content of the publication. Analysis code, and independent validation dataset supporting this review are available from the corresponding author upon reasonable request.

## Acknowledgments

The authors thank Robert W. Williams, Lu Lu, Hao Chen, and Abraham A. Palmer for providing animals, and Shelby Graham, Kyle Freeman, Sophie Pilkinton, and William Edwards for their contributions to optic nerve tissue preparation, histological processing, and manual annotations used in the independent validation datasets. We also thank Noah Emmertand Rayen Naik for their assistance with literature screening, study selection, and verification during the systematic review process.

