## Supplementary Materials for "A Systematic Review and Independent Benchmarking of Automated Nerve Morphometry Methods"

### Supplementary Figures

Supplementary Figure S1. Mean Absolute Percent Error (MAPE) Comparison Across Tools. Lower MAPE indicates better performance.

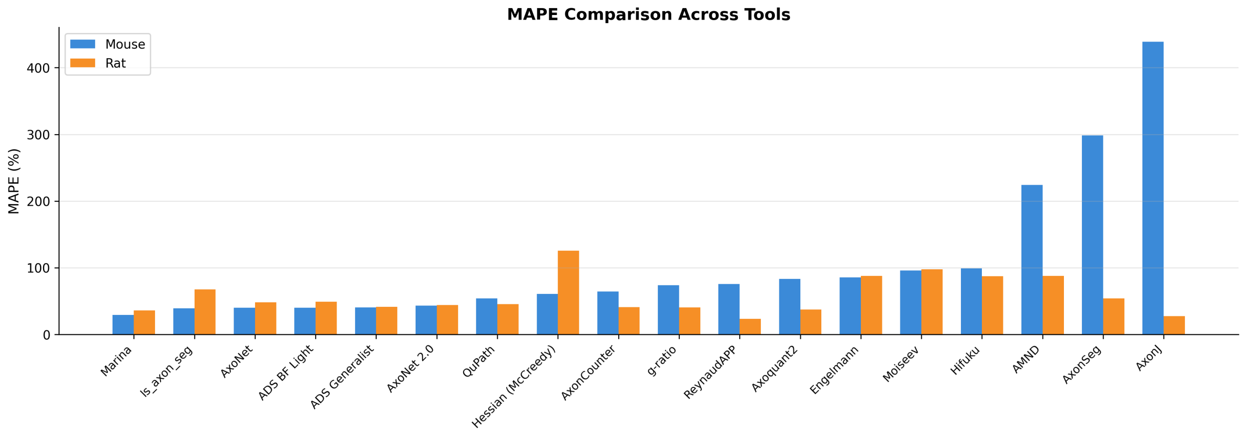


Supplementary Figure S2. Tool Performance Heatmap. Performance for each tool using paraphenylenediamine (PPD)-stained mouse and rat optic nerve images is shown. Cell colors are scaled within each metric, with green indicating better performance and red indicating worse performance. For error metrics mean absolute percentage error (MAPE) and mean absolute error (MAE), lower values correspond to better performance, whereas for Pearson correlation (r), higher values correspond to better performance. Median predicted to ground truth ratios are colored according to proximity to 1.0, with values closer to 1.0 indicating lower systematic bias.

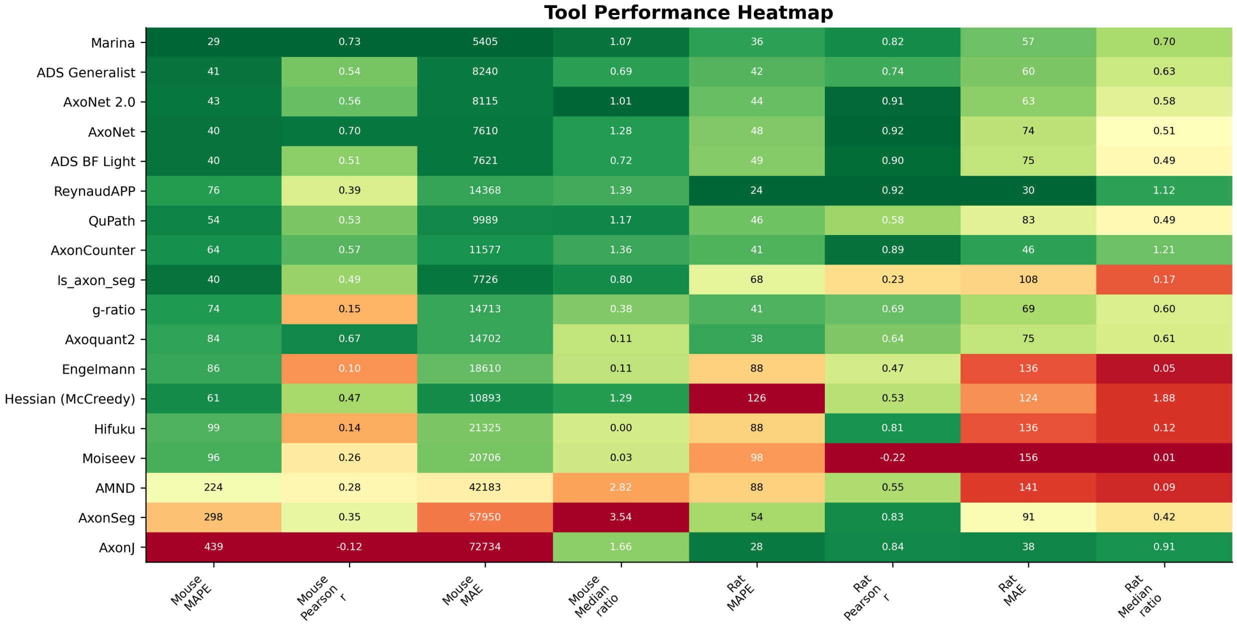

Supplementary Figure S3. The median of predicted axon count numbers from each semi- and fully automated tool were compared to the median of ground truth (GT) axon count that was manually annotated for paraphenylenediamine (PPD)-stained mouse and rat optic nerve images. *Abbreviations*: CV = computer vision; DL = deep learning.

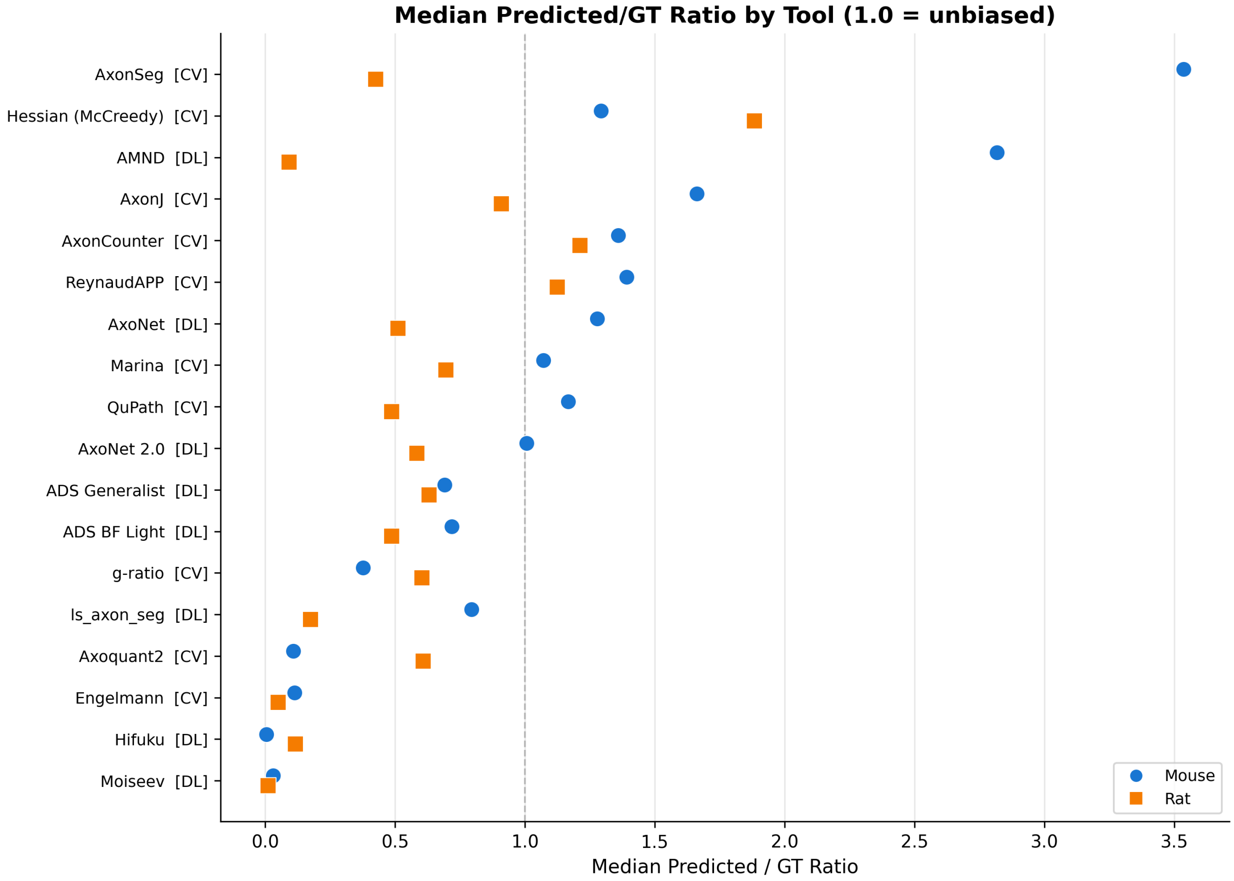

Supplementary Figure S4. Generalizability: Published vs. Independently Benchmarked Performance. Two-panel comparison showing (A) Pearson correlation r comparison between published data (diamonds) versus independently benchmarked mouse (open circles) and rat (filled squares) datasets, and (B) Dice coefficient for segmentation comparison between published data versus rat dataset benchmarked results. Delta annotations represent the performance gap.

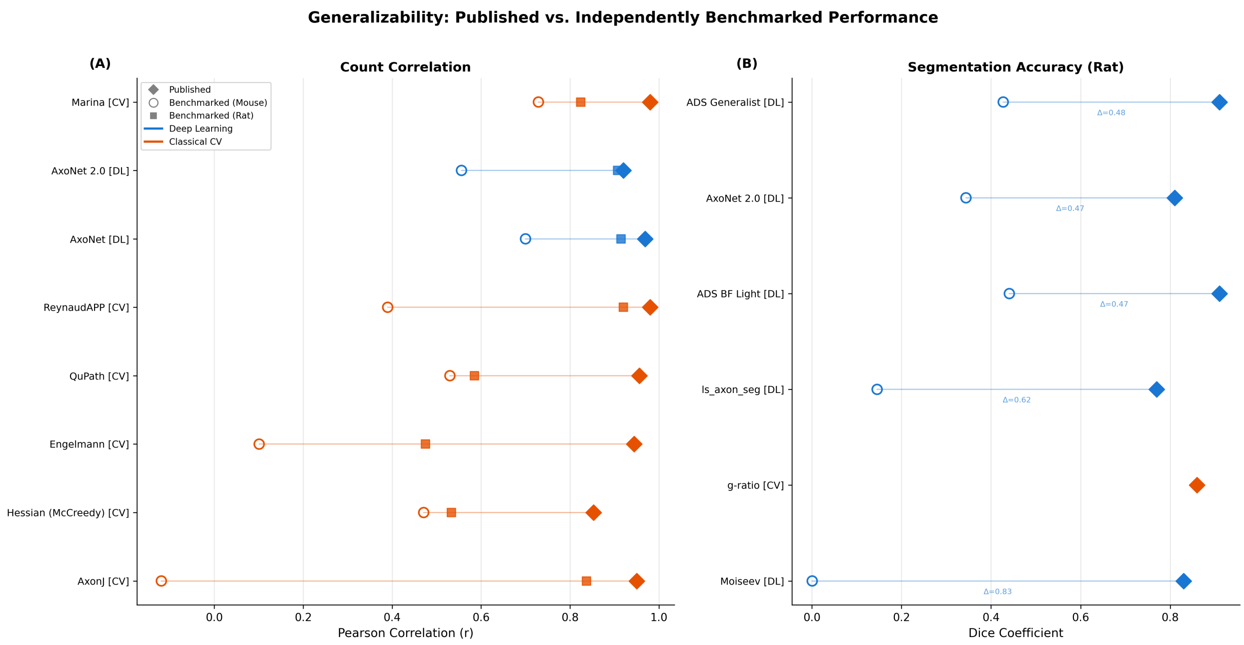

Supplementary Figure S5. Tool capability matrix summarizing the primary algorithmic approach and reported outputs for each tool. Although Moiseev enables indirect estimation of axon counts through myelin segmentation, axon counting was not reported as a primary output. *Abbreviation:* CV = computer vision.

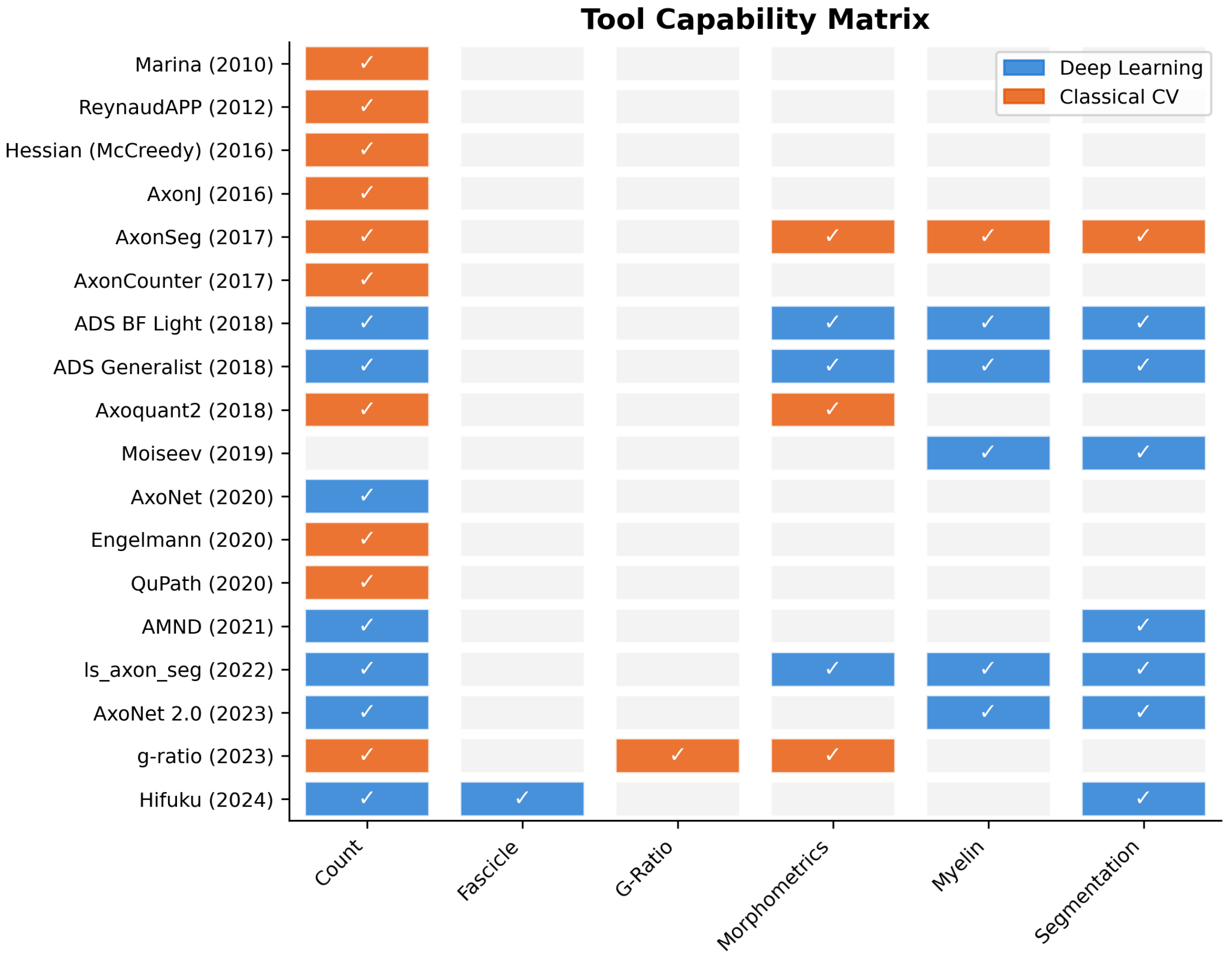

**Supplementary Tables**

Supplementary Table S1. Database Search Strategies

| Database | Search Strategy |
|---|---|
| PubMed | ("optic nerve"[Title/Abstract] OR "optic nerve head"[Title/Abstract] OR "central nervous system"[Title/Abstract] OR "white matter"[Title/Abstract] OR "nerve cross-section*"[Title/Abstract] OR axon*[Title/Abstract] OR myelin[Title/Abstract] OR "retinal ganglion cell*"[Title/Abstract] OR astrocyte*[Title/Abstract] OR microglia*[Title/Abstract] OR oligodendrocyte*[Title/Abstract] OR glia[Title/Abstract]) AND (grading[Title/Abstract] OR scoring[Title/Abstract] OR "semi-quantitative"[Title/Abstract] OR stereology[Title/Abstract] OR morphometry[Title/Abstract] OR histomorphometry[Title/Abstract] OR "axon count*"[Title/Abstract] OR quantification[Title/Abstract] OR measurement[Title/Abstract]) AND (automated[Title/Abstract] OR "computer-assisted"[Title/Abstract] OR "image analysis"[Title/Abstract] OR "digital pathology"[Title/Abstract] OR segmentation[Title/Abstract] OR algorithm*[Title/Abstract] OR "rule-based"[Title/Abstract] OR ImageJ[Title/Abstract] OR FIJI[Title/Abstract] OR CellProfiler[Title/Abstract] OR threshold*[Title/Abstract] OR watershed[Title/Abstract] OR "edge detection"[Title/Abstract] OR "machine learning"[Title/Abstract] OR "deep learning"[Title/Abstract] OR "convolutional neural network"[Title/Abstract] OR CNN[Title/Abstract]) AND (histology[Title/Abstract] OR histopathology[Title/Abstract] OR microscopy[Title/Abstract] OR immunohistochemistry[Title/Abstract]) |
| Embase | ('optic nerve'/exp OR 'optic nerve head'/exp OR 'central nervous system'/exp OR 'white matter'/exp OR 'nerve cross section*':ti,ab OR axon*:ti,ab OR myelin:ti,ab OR 'retinal ganglion cell*':ti,ab OR astrocyte*:ti,ab OR microglia*:ti,ab OR oligodendrocyte*:ti,ab OR glia:ti,ab) AND (grading:ti,ab OR scoring:ti,ab OR 'semi quantitative':ti,ab OR stereology:ti,ab OR morphometry:ti,ab OR histomorphometry:ti,ab OR 'axon count*':ti,ab OR quantification:ti,ab OR measurement:ti,ab) AND (automated:ti,ab OR 'computer assisted':ti,ab OR 'image analysis':ti,ab OR 'digital pathology':ti,ab OR segmentation:ti,ab OR algorithm*:ti,ab OR 'rule based':ti,ab OR ImageJ:ti,ab OR FIJI:ti,ab OR CellProfiler:ti,ab OR threshold*:ti,ab OR watershed:ti,ab OR 'edge detection':ti,ab OR 'machine learning'/exp OR 'deep learning'/exp OR 'convolutional neural network*':ti,ab OR CNN:ti,ab) AND (histology/exp OR histopathology/exp OR microscopy/exp OR immunohistochemistry/exp) |
| Scopus | (TITLE-ABS-KEY ("optic nerve" OR "optic nerve head" OR "central nervous system" OR "white matter" OR "nerve cross-section*" OR axon* OR myelin OR "retinal ganglion cell*" OR astrocyte* OR microglia* OR oligodendrocyte* OR glia)) AND (TITLE-ABS-KEY (grading OR scoring OR "semi-quantitative" OR stereology OR morphometry OR histomorphometry OR "axon count*" OR quantification OR measurement)) AND (TITLE-ABS-KEY (automated OR "computer-assisted" OR "image analysis" OR "digital pathology" OR segmentation OR algorithm* OR "rule-based" OR ImageJ OR FIJI OR CellProfiler OR threshold* OR watershed OR "edge detection" OR "machine learning" OR "deep learning" OR "convolutional neural network" OR CNN)) AND (TITLE-ABS-KEY (histology OR histopathology OR microscopy OR immunohistochemistry)) |

Searches conducted in January 2026. Date limits: 1999-2026. Language: English.

Supplementary Table S2. Inclusion and Exclusion Criteria for Systematic Review

| Category | Inclusion Criteria | Exclusion Criteria |
|---|---|---|
| Population / Sample | Studies involving HISTOLOGICAL cross-section analysis of nerve tissue only:<br>• Peripheral nerves (sciatic, vagus, facial, tibial, sural, etc.)<br>• Optic nerve<br>• Spinal cord white matter tracts<br>• Cranial nerves<br><br>Imaging modalities MUST be histology-based:<br>• Electron microscopy (TEM, SEM)<br>• Light microscopy<br>• Confocal microscopy of fixed tissue sections<br>• Histological stains (toluidine blue, osmium tetroxide, immunohistochemistry, etc.) | • Studies not involving nerve tissue cross-sections<br>• ALL in-vivo imaging studies (MRI, OCT, DTI, diffusion imaging) - even if combined with histology<br>• Brain parenchyma studies (cortex, hippocampus, striatum, cerebellum) without nerve tract cross-sections<br>• Retinal imaging or flat-mount studies without optic nerve histological cross-sections<br>• Cell culture studies without nerve tissue sections<br>• Studies focused solely on non-neural structures<br>• Live imaging or intravital microscopy |
| Intervention / Method | Automated or semi-automated computational tools for HISTOLOGICAL nerve cross-section quantification:<br>• Machine learning / deep learning segmentation tools (e.g., AxonDeepSeg, AxoNet, U-Net based methods)<br>• Classical image analysis pipelines (ImageJ/FIJI macros, CellProfiler, QuPath, ilastik)<br>• Custom algorithms for axon/myelin segmentation from histological images<br>• Tools measuring: axon counts, fiber diameter, myelin thickness, g-ratio, axon density, fiber size distribution<br><br>Tools must be applied to histological tissue sections, not in-vivo images. | • Studies with no automated/computational quantification component<br>• Manual-only counting or grading without tool development/validation<br>• ALL MRI-based analysis tools (tractography, diffusion tensor imaging, myelin water imaging)<br>• ALL OCT-based analysis tools (including OCT angiography)<br>• ALL in-vivo imaging analysis methods<br>• Glial cell quantification tools not applied to nerve cross-sections<br>• Studies applying tools only to brain tissue or non-nerve structures<br>• Electrophysiology, behavioral, or functional studies without histological analysis |
| Outcomes | Studies that report:<br>• Performance metrics for the quantification tool: accuracy, precision, recall, sensitivity, specificity, Dice score, Jaccard index, F1-score, IoU<br>• Validation metrics: inter-rater reliability, | • Studies without quantitative outcomes related to nerve morphometry<br>• Studies that use a tool but report NO performance metrics or validation<br>• Studies reporting only biological findings without evaluating the quantification |

| | | |
|---|---|---|
| | intra-class correlation, coefficient of variation, Bland-Altman analysis<br>• Comparison to ground truth or gold standard (manual segmentation of histological images)<br>• Quantitative morphometric outputs with method validation: axon counts, myelin thickness, g-ratio, fiber diameter distributions<br><br>Tool must be validated on histological images. | method<br>• Studies without any comparison to ground truth, manual counts, or reliability assessment<br>• Purely descriptive studies without morphometric quantification<br>• Validation performed only on in-vivo imaging data |
| Study Type | • Original research articles (basic science, translational, preclinical, or clinical)<br>• Methods papers describing new histological analysis tools or pipelines<br>• Validation studies comparing quantification approaches on histological images<br>• Tool development papers with performance evaluation on nerve histology | • Reviews, editorials, commentaries<br>• Conference abstracts without full methodology/data<br>• Non-peer-reviewed gray literature<br>• Preprints without sufficient methodological detail<br>• Case reports without tool validation<br>• Studies focused solely on in-vivo imaging methodology |
| Language | English. | Non-English publications. |

**Supplementary Table S3. Extracted dataset from 71 included studies.**

https://docs.google.com/spreadsheets/d/1GUSp4tOeOHJblubaowRdC1MAOHv4gbs2/edit?usp=sharing&ouid=111131618220983953558&rtpof=true&sd=true
